# Improving visual attention following right hemisphere stroke: A preliminary study

**DOI:** 10.1101/2022.08.19.504424

**Authors:** Grace Edwards, Laurel J. Buxbaum, Gang Chen, Dylan Edwards, Lorella Battelli

**Affiliations:** Center for Neuroscience and Cognitive Systems@UniTn, Istituto Italiano di Tecnologia, Rovereto, Italy; Department of Psychology, Harvard University, Cambridge, MA 02138, USA; Moss Rehabilitation Research Institute, 50 Township Line Rd, Elkins Park, PA 19027, USA; Department of Rehabilitation Medicine, Thomas Jefferson University, 901 Walnut Street, Philadelphia, PA 19107; National Institute of Mental Health, Bethesda, MD, 20892, USA; Berenson-Allen Center for Noninvasive Brain Stimulation and Department of Neurology, Beth Israel Deaconess Medical Center, Harvard Medical School, Boston, Massachusetts, USA, 02215

**Keywords:** Spatial attention, Sustained attention, right hemisphere cerebrovascular accident, lateralized attention, attention rebound

## Abstract

Left inattention is common in individuals following right cerebrovascular accident (RCVA). In neurotypical adults, we have previously found prolonged rightward visual attention resulted in a subsequent increase in leftward attention. Here we applied the same method in neurological patients with RCVA and found improved post-intervention attention both to the left and right of visual fixation in participants with mild to no leftward inattention in comparison to a control. No such benefit was detected in participants with more pronounced leftward inattention. Given the feasibility of the intervention which leverages performance in the right unaffected visual space, future studies should examine the longevity and generalizability of such an intervention to other attention demanding tasks.

## Introduction

Globally, over one in four people will experience a stroke in their lifetime (Lindsay et al., 2019). Stroke is the most common cause of unilateral brain damage, among traumatic brain injury, disease, seizures, and infections (Silasi & Murphy, 2014). One symptom of right cerebrovascular accident (RCVA) is leftward inattention. Moderate and severe leftward inattention, termed hemispatial neglect, can occur in 23-26% of cases and is typically diagnosed in the acute phase after a stroke (Ogden, 1985; Becker & Karnath, 2007). The prognosis following initial diagnosis of leftward inattention varies, including full recovery with rehabilitation (Apperlros et al., 2004). While in most cases mild forms of inattention go undetected with standard neuropsychological measures, they are an underlying deficit that may prevent full recovery of everyday activities. Here, we present preliminary findings which suggest increased sustained attention in patients with RCVA.

The cause of leftward inattention is likely twofold. First, left lateralized attention is impacted directly by damage to the right hemisphere (Kinsbourne 1987; Batholomeo & Malkinson, 2019). Lateralized attention is supported by a cortico-subcortical network including the intraparietal sulcus (IPS), frontal eye fields (FEF), superior parietal lobe (SPL), dorsolateral prefrontal cortex (DLPFC), the superior colliculus, pulvinar nucleus, and cholinergic inputs from the basal forebrain (Corbetta & Shulman, 2002; Shipp, 2004; Martinez & Sarter, 2004; Buschman & Miller, 2007). Damage to these regions in the right hemisphere can significantly impact leftward attention. Second, due to the right hemispheric damage, a subsequent imbalance in activity between left and right lateralized attention processing regions may further impact leftward attention. Typically, left and right lateralized attention processing regions mutually inhibit each other to enable equal lateralized attention. With right hemisphere damage, inhibition from right to left may be reduced, resulting in hyperactivity in the left hemisphere, and therefore further inhibition of the stroke-affected regions (Kinsbourne, 1993; Kinsbourne, 1994; Corbetta et al., 2005).

Significant effort has culminated in multiple rehabilitation methods, which vary in complexity and success. Some rehabilitation methods have focused on compensatory mechanisms, such as Scan Therapy (Parton et al., 2004), where participants are trained to shift their gaze to the left visual field thereby continuously moving information into their putatively unimpaired right visual field. However, the generalizability of scan therapy is limited outside of training (Kerkhoff et al., 2021) and is not aimed at recovery of leftward inattention, but rather a strategy that might help overcome the deficit. The deficit remains detectable when subjects are presented with bilateral visual field tasks, a condition arguably more similar to natural viewing. The application of prismatic goggles has been more successful (Rossetti et al., 1998; Frassinetti et al., 2002). The goggles shift the perceived location of the body and objects to the right, and continuous motor training with the adapted visual input results in an adaptation after-effect in which attention and action are left-shifted. These rehabilitation methods often don’t address recovery within the impaired hemispace, while other methods have focused on regaining leftward attention. These methods include training with centrally presented cues to promote leftward attention (Làdavas et al., 1994) and rebalancing the right and left hemispheric activity with inhibitory noninvasive brain stimulation to the healthy hemisphere to distribute visual attention across the entirety of space (Brighina et al., 2003; Agosta et al., 2014). These rehabilitation methods either involve complex technology, or the participants’ capacity to maintain attention to the affected visual field (Barrett et al., 2006).

Here, we present preliminary results on a potential new leftward attention-enhancing method which provides “proof of concept” for future development of rehabilitation interventions. In neurotypical individuals, we previously found 30 minutes of sustained unilateral attention resulted in an increase in attention in the opposite visual space (Edwards et al., 2021). It is plausible the attention increase was caused by an overshoot in activity as lateralized attention processing regions rebalance following prolonged imbalance (Edwards et al., 2021). When attention is directed to one side of visual space with a centrally held fixation for a prolonged period, attention to the other visual space may be actively suppressed. When returning to whole field attention, sustained suppression of the other visual space is released, potentially resulting in a post-inhibitory rebound (Pugh & Raman, 2006), or a homeostatic gain control response (Muret & Makin, 2021; Turrigiano, 2017). The visual space-specific attention increase shows potential for application in patients with leftward inattention. Similarly, here participants with RCVA were required to fixate centrally and attend rightward, toward their unimpaired visual space, to perform a demanding sustained attention task. We predicted an improvement in leftward attention, as was previously found in neurotypical individuals (Edwards et al., 2021). We expected that patients with less leftward inattention would experience a greater impact of intervention, as severity of leftward inattention tends to negatively correlate with successful intervention (Gillen et al., 2005). Furthermore, it is also plausible that those with less leftward inattention maybe be experiencing less ongoing intercortical inhibition from the healthy hemisphere (Brighina et al., 2003; Agosta et al., 2014), therefore might be more permissive of an intervention response.

## Methods

### Participants

Fourteen participants with chronic (> 6 months post stroke onset) right hemisphere cerebrovascular accident (RCVA) were recruited from the Neuro-Cognitive Research Registry at Moss Rehabilitation Research Institute. Exclusion criteria included: Visual field cuts, second stroke, active bleeding, major organ failure, complex or terminal medical illness, major surgery, previous neurologic or psychiatric condition, coma or severe cognitive impairment, and radiological evidence of nonvascular cause. Importantly, participants were not selected due to reported inattention. We planned to characterize leftward inattention, and use the outcome as a predictor of the proposed intervention impact (see *Characterizing contralesional attention*). Three participants recruited were excluded prior to data collection due to undocumented visual field cuts determined via a manual visual perimetry task (see *Procedure*). Table 1 reports participant demographics and scores on the National Institutes of Health Stroke Scale. Table 2 reports the radiology summary of each participant collected during the acute stroke hospital admission. All participants gave written informed consent in accordance with the Institutional Review Board at the Albert Einstein Healthcare Network.

**Table 1:**
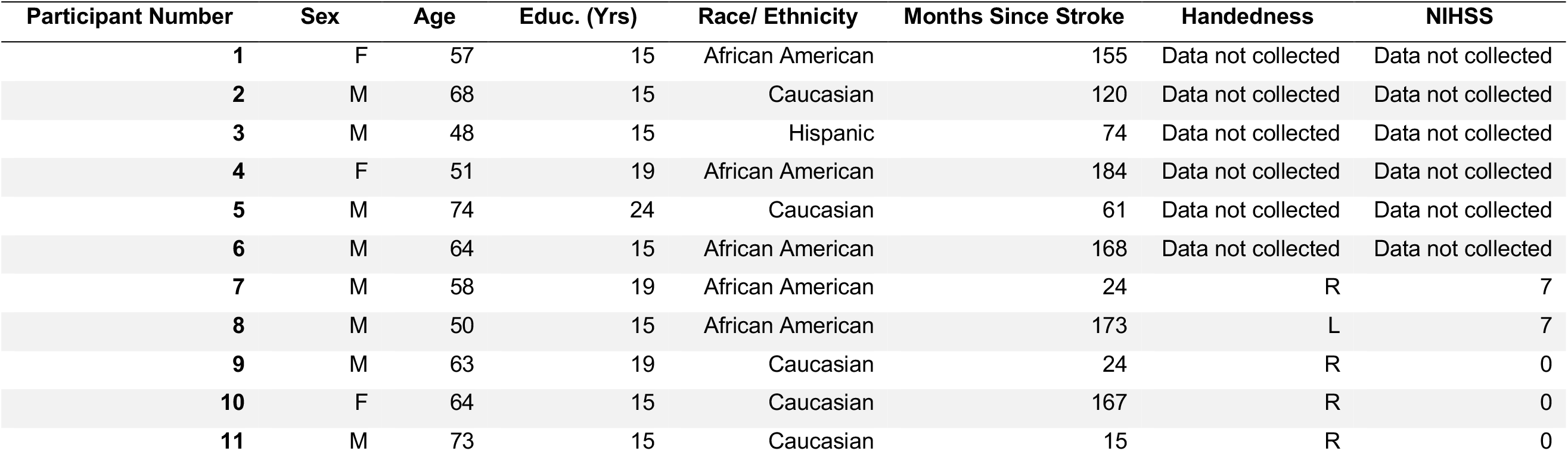
Participant demographics and neuropsychological characteristics. The National Institute of Health Stroke Scale (NIHSS) indicates the severity of the stroke measured at the time of the study. Cognitive functions such as language, attention, perception, and level of consciousness are measured at bedside. When collected during the acute phase of the infarction, the severity of the stroke is classified as follows: 0 = no stroke symptoms, 1–4 = minor stroke, 5–15 = moderate stroke, 15–20 = moderate/severe stroke and 21–42 = severe stroke.

**Table 2:** Summary of radiology results from acute phase of infarction.

| Participant Number | Scan Type | Radiology summary |
| --- | --- | --- |
| 1 | CT-Clinical | Hemorrhage right frontal intracerebral hemorrhage |
| 2 | CT-Clinical | Ischemic stroke basal ganglia right |
| 3 | Unknown-Clinical | Ischemic stroke basal ganglia right |
| 4 | CT-Clinical | Hemorrhage basal ganglia right |
| 5 | MRI-Clinical | Ischemic stroke unspecified right, acute infarct in right subcortical region |
| 6 | MRI-Clinical | Ischemic stroke occipital right; Ischemic stroke basal ganglia left-small lacunar infarct in left caudate head |
| 7 | CT-Clinical | Large transcortical infarct involving nearly the entire right middle cerebral artery territory. |
| 8 | CT-Clinical | Ischemic stroke middle cerebral artery right |
| 9 | MRI-Clinical | Ischemic stroke middle cerebral artery right |
| 10 | MRI-Clinical | Hemorrhage frontal right |
| 11 | CT-Clinical | Large right middle cerebral artery stroke |

### Stimuli

#### Bilateral Multiple Object Tracking (MOT)

Bilateral MOT involves tracking objects among distractors on either side of fixation (Yantis, 1992). Bilateral MOT has been demonstrated to recruit bilateral attention processing regions (Pylyshyn & Storm, 1988; Culham et al., 1998). Each trial begun with four black dots (radius 0.25°) presented on either side of a central fixation point for 1000 ms (Figure 1a). Eye-tracking data were recorded throughout the session (*see Eye-Tracking Acquisition and Analysis*) Two dots on either side of fixation flashed to indicate that they were the targets participants were to covertly track. The dots then moved among the distractor dots for 4000 ms. All dots remained within their original space (to the left or right of fixation), never crossing the vertical meridian. The dots moved at a constant speed (degrees per second), determined by each participant’s individual threshold (see *Thresholding* and *Procedure*). During the movement period, the dots repelled one another to maintain a minimum of 1.5° space. After 4000 ms the dots stopped and one dot either to the left or right of fixation was highlighted in red. The participants were asked to respond if the red dot was a “target” or a “distractor” by a button press. Importantly, the participants were unaware of which side of fixation would be tested, necessitating sustained covert attention to both the left and right visual space during the movement period. After the participant responded, the central fixation point turned green to indicate that they were correct or red to indicate that they were incorrect.

**Figure 1:**
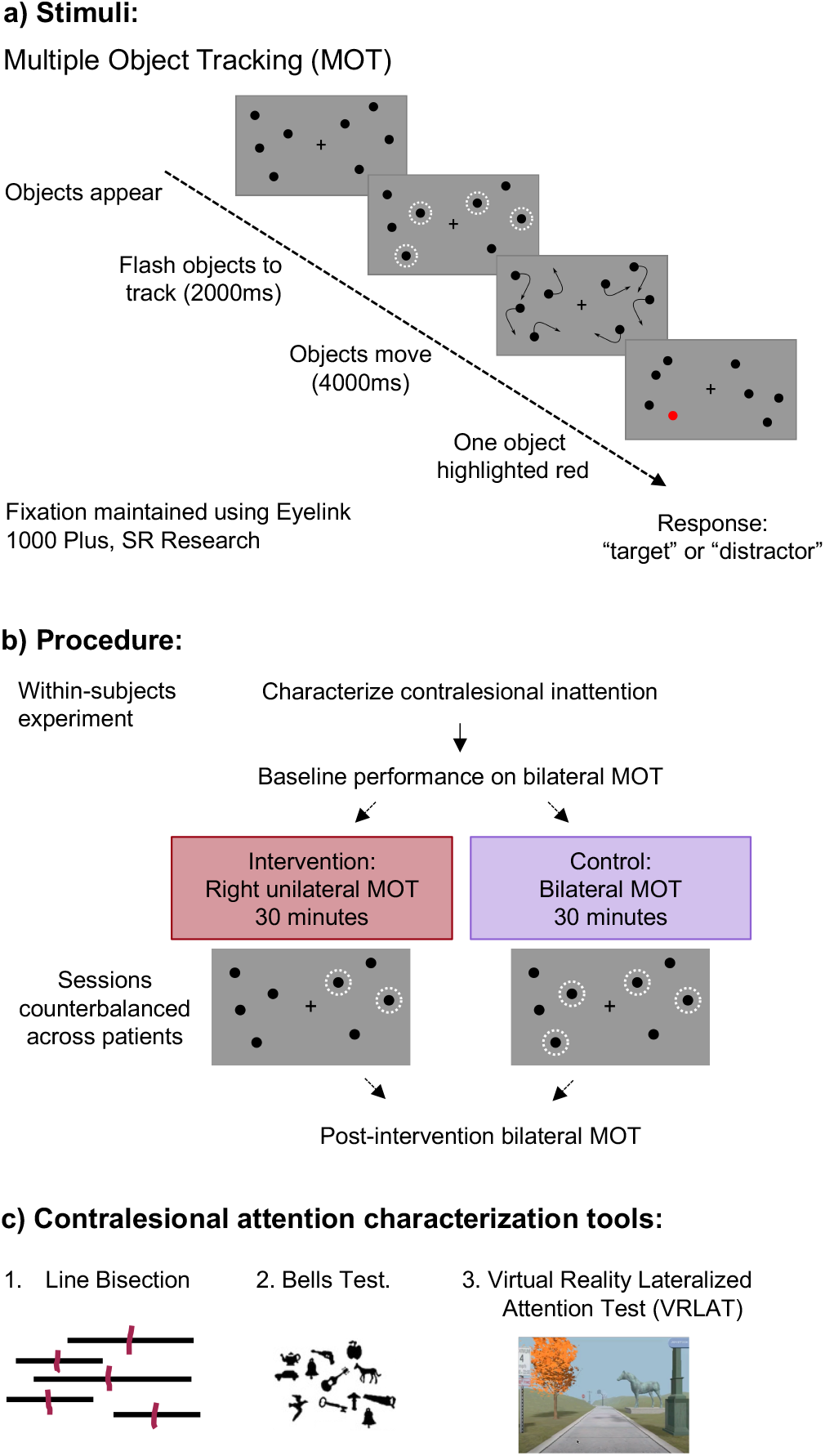
Methods and procedure. 1a) depiction of one trial of bilateral multiple object tracking. 1b) Two sessions were counterbalanced across participants. Each session was the same, except for the intervention (in red) and control (in purple) periods. 1c) Attention characterization tools: i) Line Bisection (17 lines), ii) Bells Test (35 bells in 265 distractor objects), iii) Virtual Reality Lateralized Attention Test (VRLAT; name color of trees and animal statues presented left and right of path).

#### Unilateral Multiple Object Tracking

Unilateral MOT was employed to isolate attention to the (ipsilesional) right of fixation during the intervention (Edwards et al., 2021).

Similarly to bilateral MOT, each trial began with four black dots presented on either side of fixation. Then two dots to the right of fixation flashed to indicate that these were the targets to track for the duration of the movement period. Importantly, participants were only tracking dots to the right of fixation, however, four dots were still present to the left of fixation to equalize sensory input across both visual spaces. With this manipulation, we only modulated lateralized visual attention, rather than lateralized processing of visual input. After 4000 ms, the dots stopped moving, and one dot to the right of fixation was highlighted in red. Participants responded if that dot was a “target” or a “distractor” via a button press and received feedback on their performance.

#### Thresholding multiple object tracking speed

Participants’ performance was set at 75% correct at the beginning of each session by modulating the speed of the moving objects. Testing every patient at their individually-set performance threshold, we ensured the tasks were equated for difficulty across patients and tasks, and that any improvement or deterioration was due to the intervention. All participants begun with 16 practice trials with speed set at 2° per second. In the following five blocks of thresholding, the speed of the objects were randomly assigned from 2° to 16° per second with 16 trials per block. Linear interpolation determined the speed at which each participant performed at 75% correct.

#### Characterizing contralesional attention

We characterized the presence and extent of contralesional attentional deficit using three neuropsychological measures: 1) Line Bisection task (Schenkenberg et al., 1980; Bisiach et al., 1983; Wilson et al., 1987), 2) Bells Test (Gauthier et al., 1989), and 3) Virtual Reality Lateralized Attention Test (VRLAT, Buxbaum et al., 2012). The Line Bisection task is a pencil and paper task requiring participants to bisect 17 horizontal lines (see Appendix 1). The experimenter (author GE) demonstrated the line bisection on one line on a separate test sheet, and then gave the participant their own sheet to perform the bisections. The sheet was held square in front of the participant until they finished bisecting all 17 lines. Participants with leftward visual attention deficit bisect the lines more than 0.6 cm rightward of midpoint (Zeltzer & Menon, 2008). Neurotypical controls have been shown to have a slight leftward bias, but only to ∼0.15 cm to the left of the midpoint (Cavézian et al., 2008; Benwell et al., 2014; Learmonth et al., 2018), and often with large variance across the sample (Learmonth & Papadatou-Pastou, 2021). Theoretically, those with leftward attentional deficits do not attend to the left end of the line, resulting in a rightward perception of the midpoint of the line. The Bells Test is a pencil and paper task requiring patients to circle bells embedded in 265 distractors (see Appendix 1). There were 35 bells spread evenly across a landscape A4 sheet of paper. Seventeen bells were presented to the left of center, 17 to the right, and one in the center. Detection of 11 or fewer bells (omission of more than 6) on the left of the sheet of paper was considered the cut-off to diagnose leftward visual attention deficit based on standardized measures collected on age-matched controls (Gauthier et al., 1989). The VRLAT was presented on a 15.5 × 27.5-inch flat screen, 34 inches from the participant (Buxbaum et al., 2012). The VRLAT was run at the enhanced-level on a PC with an Intel core 2 Duo processor.

The participants first practiced driving down a pathway in a virtual reality environment, navigating with a Logitech Attack 3 joystick. Once comfortable with the operation, the participants begun the task: while navigating along a winding path, the participants were asked to name the color of the trees and the identity of animal statues they passed either to the left or right of the path. The participants were asked to ignore stationary and moving distractor objects, and to ignore auditory distractors such as the sound of a car engine. The participants drove in both directions along the path, so all the targets were viewed once on the left of the path, and once on the right. Detection of 18 or less targets on the left of the path indicated likely leftward inattention in comparison to age-matched controls (Buxbaum et al., 2012; see Appendix 1). Participants who were below the clinical cut off on one or more characterization tests were determined to have leftward inattention (n=6; in grey Table 3). Those above the clinical cut-off were classified as having mild to no leftward inattention (n=5; Table 2).

**Table 3:**
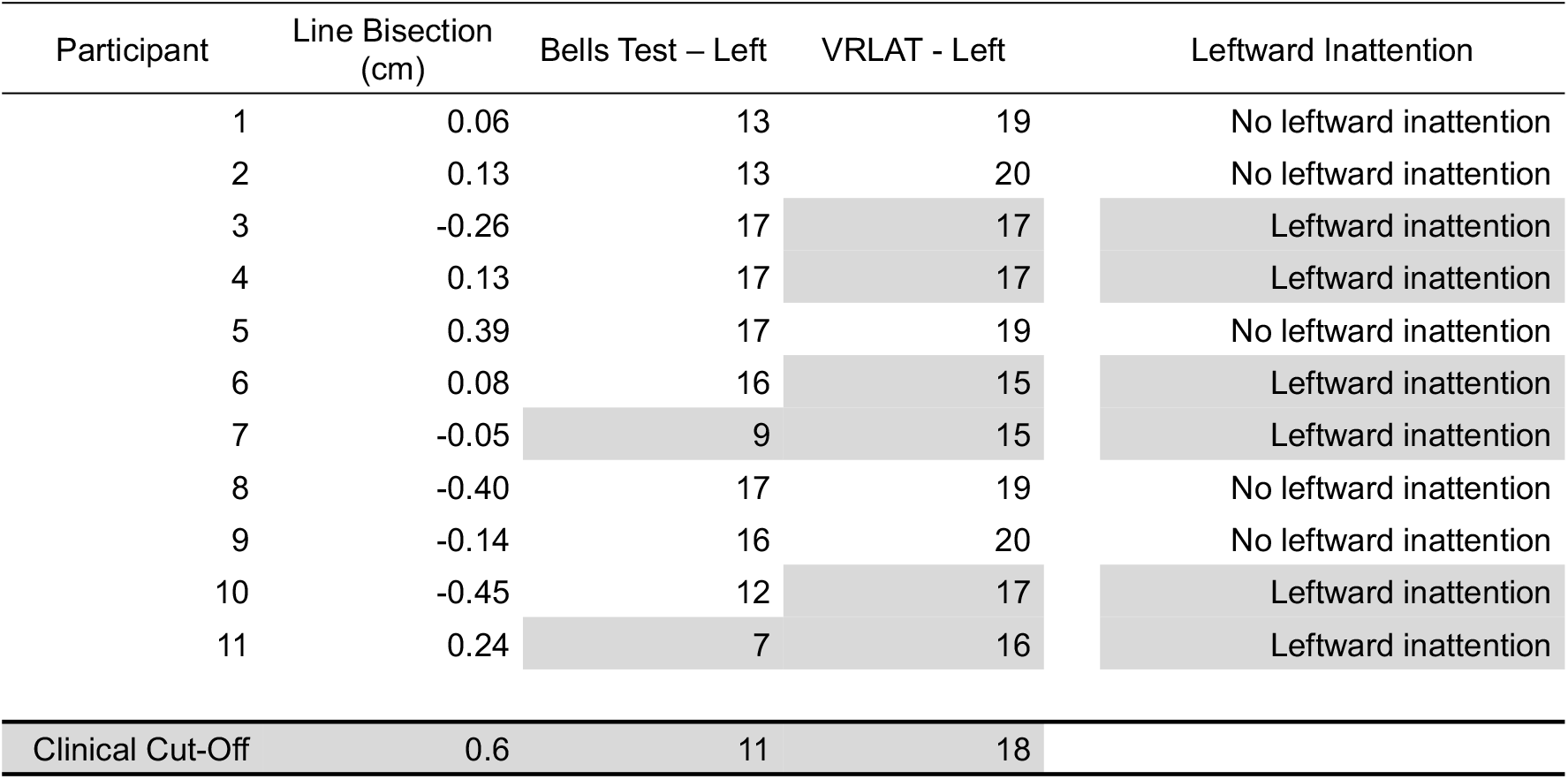
Attention characterization scores. Line bisection, Bells Test, and VRLAT scores presented for each participant. The clinical cut-off scores are presented in the final row. Individuals with a score below the clinical cut-off in any of the three tests were grouped into those with leftward inattention (highlighted in grey).

#### Procedure

Participants were tested on two separate days, each session lasting a maximum of 2 hours (Figure 1b). In one session, participants performed the intervention task, in the other the participants performed a control task. These sessions were counterbalanced across participants to control for cross-over effects. In the first session, participants were tested for visual extinction and visual field cuts using a manual extinction and visual perimetry test. In both sessions, the participants performed contralesional attention characterization using Line Bisection, Bells Test, and Virtual Reality Lateralized Attention Test (VRLAT; see *Characterizing contralesional attention*). Next, we thresholded the speed of the dots in the MOT so each participant was performing at 75% correct prior to any intervention (see *Thresholding multiple object tracking speed*). After thresholding, the participants performed 10 minutes of bilateral MOT to determine their baseline performance prior to the intervention or control. Next on the intervention day, the participants performed right unilateral MOT for 30 minutes. Alternatively, next on the control day, participants performed bilateral MOT for 30 minutes. Participants were allowed to take breaks during the manipulation phase. After the intervention or control tasks, participants performed 10 minutes of bilateral MOT to determine if their performance had been modulated to the left or right of fixation by the intervention or control task.

#### Eye-tracking acquisition & analysis

An Eyelink 1000 Plus eye-tracking system (SR Research) was used to record participants fixation location during each trial of the experiment (Supplemental Figure 1). Each run began with a calibration and validation of the location of the pupil on the screen. If the drift correction failed, the participants would be re-calibrated. Participants could blink comfortably without restarting the trial. Once the data were collected, the percentage of time participants maintained a central fixation (within a 3° x 3° boundary box of fixation) during the trial was used as a predictor in the linear mixed effects model. Eye-tracking data from three subjects per session could not be collected due to equipment issues. Linear mixed effects modeling uses maximum likelihood estimation to cope with the missing data-points.

#### Behavioral Data analysis

A general linear mixed effects model was fit to the participants’ multiple object tracking accuracy in R (R Core Team, 2019) using *glmer()* function from the *lme4* package (Bates et al., 2014). The between-subjects predictors were the presence of leftward inattention (scored categorically, see Table 3) and percentage of time participants maintained central fixation during the manipulation period. The three within-subject predictors were manipulation (Intervention or Control), Session (pre-or post-intervention), and visual space (Left or Right of fixation). The output and model comparisons indicated no difference between the left and right of fixation (see Supplemental information for comparisons), so we collapsed across space, essentially averaging performance on left and right MOT. Following model comparisons, the selected model was: *glmer (Accuracy ∼ Session * Manipulation * InattentionGrouping + Eyetracking* + *(1+Session*|*Subs) + (1+ Manipulation* |*Subs), data = AttIso, family = binomial)*. The linear mixed effects model included random intercepts and slopes. Interactions and main effects are reported with *chi-squared* and *p-values*. Individual contrasts were performed using *emmeans()* and *adjust=“mvt”* for multiple comparisons (Lenth et al., 2020). Data and analysis scripts are available to reviewers and will be available to all on OSF when published.

## Results

We found a main effect of leftward attention group (left inattention versus no left inattention; χ^2^(1)=16.5781, p<0.0001, *glmer*), no statistical evidence for the main effect of session (pre-versus post-manipulation; pre-versus post-intervention; χ^2^(1)=0.4959, p=0.4813, *glmer*), no statistical evidence for the main effect of manipulation (intervention versus control task; χ^2^(1)=0.0328, p=0.8563, *glmer*), and no statistical evidence for the main effect of visual space (left versus right; χ^2^(1)=0.0713, p=0.7895, *glmer*). We also found no statistical evidence for a four-way interaction between left attention group x session x manipulation x visual space (χ^2^(1)=0.1603, p=0.6889, *glmer*), however, our model output indicated strong statistical evidence for a significant three-way interaction for left attention group x session x manipulation, collapsing across visual space (χ^2^(1)=7.3810, p=0.0066, *glmer*). Collapsing across visual space was also supported as the best fit for our data when performing model comparisons (see Supplemental Information).

Collapsing across visual space, in the group of participants with mild to no leftward inattention (n=5), we found a significant difference in MOT accuracy between the intervention and control tasks at the post-manipulation time point (Figure 2 & 3a; estimate=0.565, se=0.226, z=2.503, p=0.0245, confidence interval (CI)=0.0602, 1.069; *emmeans()* with *adjust “mvt”*). However, we found no statistical evidence for the difference from pre-to post-intervention (estimate=0.3228, se=0.230, z=1.401, p=0.5012, confidence interval (CI)=-0.251, 0.896; *emmeans()* with *adjust “mvt”*) or from pre-to post-control task (estimate=-0.4394, se=0.253, z=-1.735, p=0.2900, confidence interval (CI)=-1.070, 0.191; *emmeans()* with *adjust “mvt”*). Nevertheless, these results suggest that at the post-manipulation timepoint, there was greater MOT accuracy to both the left and right of fixation with the intervention in comparison to the control task.

**Figure 2:**
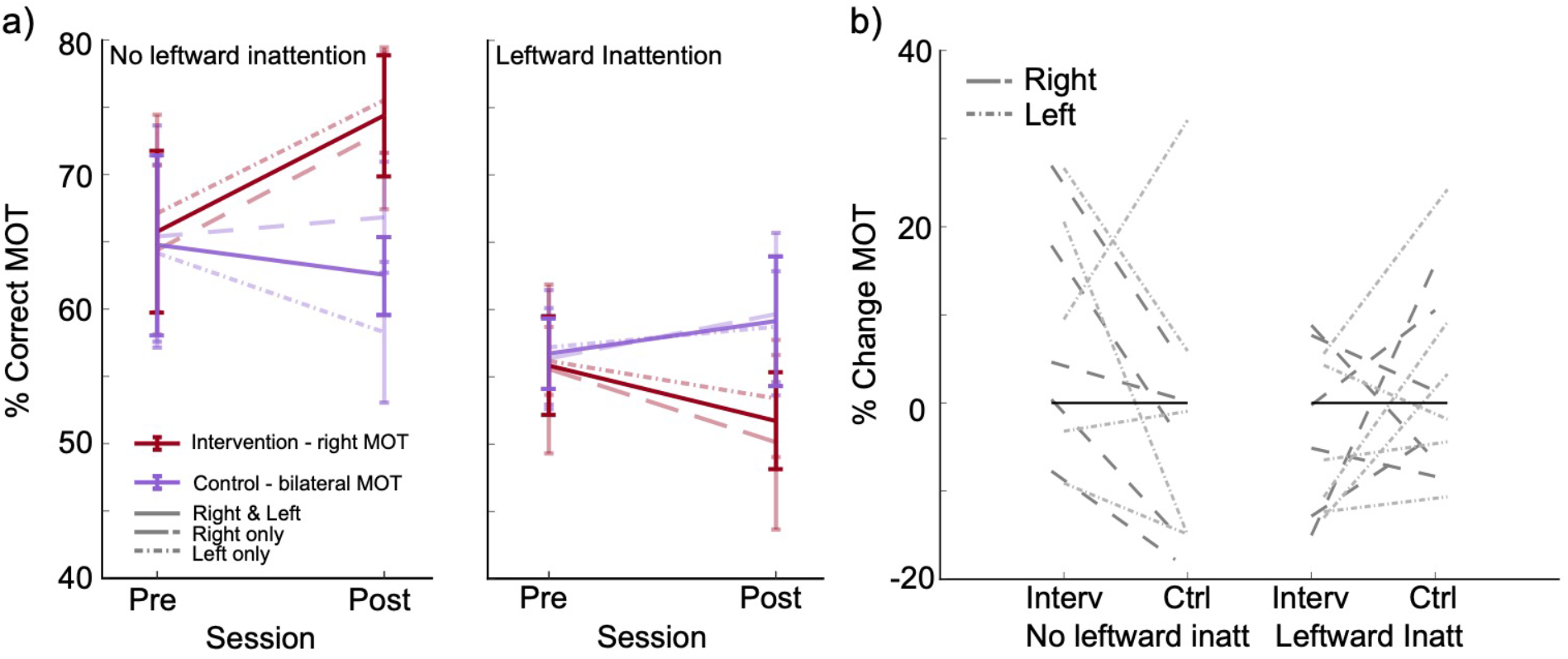
Raw percentage correct multiple object tracking. a) Pre- and post-intervention and control sessions for group with no leftward inattention (LEFT) and with leftward inattention (RIGHT). b) Individual subject percentage change between pre-to post-intervention and control sessions for group with no leftward inattention (LEFT) and with leftward inattention (RIGHT).

In contrast, the participants with leftward inattention (n=6) demonstrated no post-manipulation impact of intervention when compared with control (Figure 2; estimate=-0.380, se=0.190, z=1.999, p=0.0891, confidence interval (CI)=-0.0451, 0.8046; *emmeans()* with *adjust “mvt”*). Likewise, no statistical evidence was found for the difference from pre-to post-manipulation for the intervention (estimate=-0.3637, se=0.203, z=-1.788, p=0.2619, confidence interval (CI)=-0.870, 0.143; *emmeans()* with *adjust “mvt”*), or control task (estimate=0.0123, se=0.180, z=0.068, p=1.0000, confidence interval (CI)=-0.436, 0.461; *emmeans()* with *adjust “mvt”*).

In a *post-hoc* analysis, we correlated the magnitude of the difference between the intervention and control task at the post-manipulation time point with scores from the attention characterization tests using Spearman correlations. We found that more accurate report of left-sided objects in the VRLAT was associated with relatively better attentional tracking for the intervention relative to control task (Figure 3a; rho=0.82, p=0.006 Bonferroni corrected, CI(0.34,0.92), confirming that individuals with mild or no left inattention showed a greater difference between the intervention and control at the post-manipulation time point. No such correlation was evident for right-sided performance in the VRLAT (rho=0.35, p=0.88 Bonferroni corrected, CI(-0.36, 0.77)), leftward or rightward performance in the Bells Test (Figure 3b; left: rho=0.3, p=0.73, CI(-0.41,0.74); right: rho=0.24, p=0.94, Bonferroni corrected, CI(-0.33, 0.78)), or performance in the Line Bisection (Figure 3c; rho=-0.15, p=0.94 Bonferroni corrected, CI(-0.65, 0.55)).

**Alternative Figure 3.**
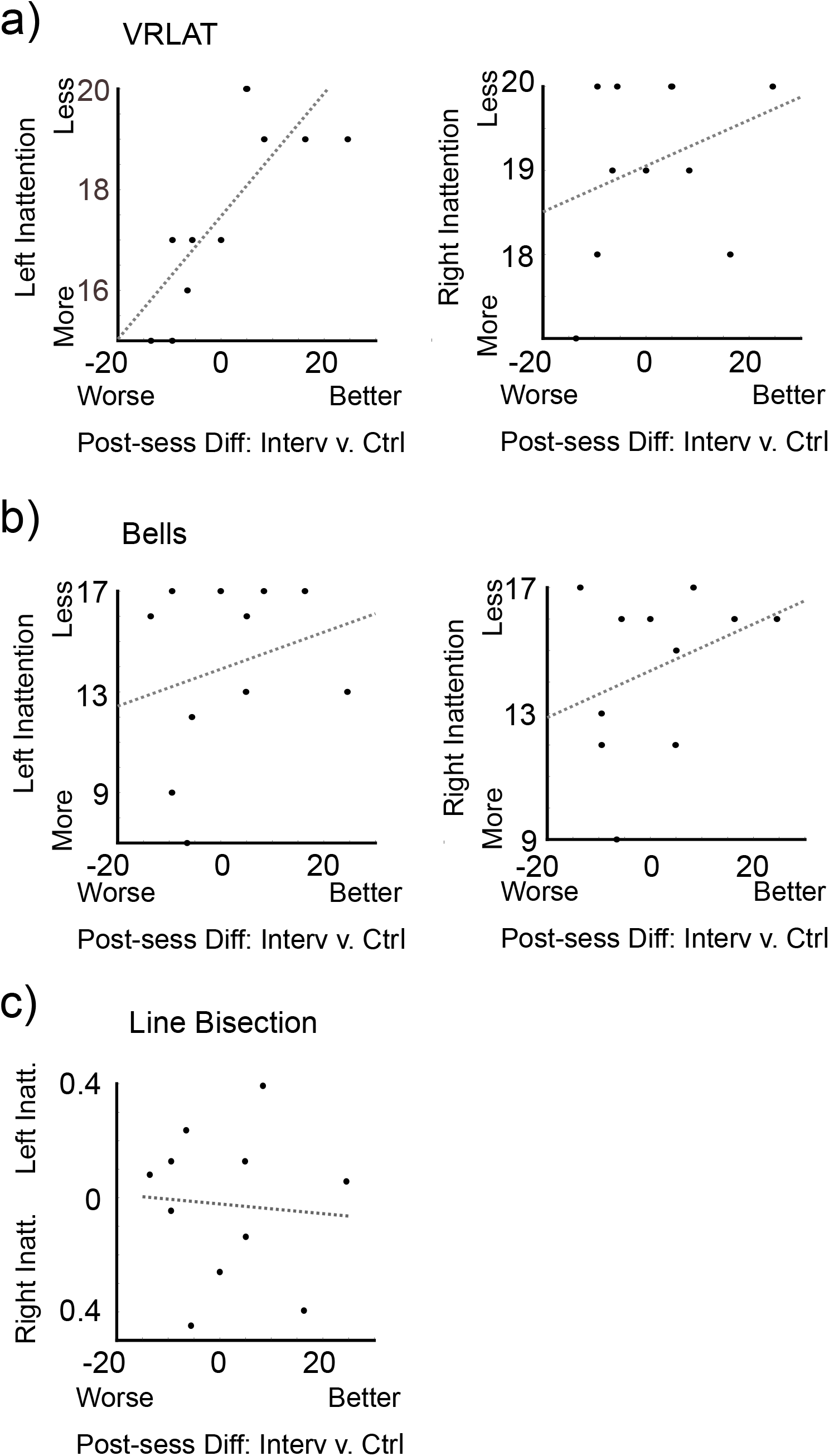
Difference between multiple object tracking accuracy performance (%) in intervention and control conditions at the post-manipulation time point correlated with attention tests. Lines depict least-squares estimates. **a)** MOT difference score plotted against Left and Right VRLAT score. **b)** MOT difference score plotted against left and right Bells Test score. **c)** MOT difference score plotted against Line Bisection score.

## Discussion

At the post-manipulation time-point, we demonstrated that prolonged attention isolation to the right of fixation was associated with better performance in attentional tracking ability as compared with a control intervention in participants with mild to no leftward inattention. Due to the lack of tracking performance change from pre-to post-intervention, our findings are limited to post-intervention sustained attention performance in comparison to the control. In the control, we found sustained bilateral attention was associated with a trending decline in performance from pre to post. With the intervention, sustained unilateral attention was associated with a comparative increase in bilateral attention performance post-intervention. These findings suggest that the intervention may facilitate sustained attention performance.

Performance for participants with more severe leftward inattention was not affected by the attentional manipulation. The distinction in impact of intervention for those with and without leftward inattention is not uncommon in studies of post-stroke recovery (Gillan et al., 2005; Winstein et al., 2016). Those with greater leftward inattention tend to progress more slowly than those who have less leftward inattention. The relatively better performance across the visual array post-intervention as compared to post-control for those with mild to no leftward inattention indicates this simple, yet effective, intervention may be at least temporarily beneficial for those with RCVA. This procedure capitalizes on a task that individuals with RCVA can perform easily – a task which is presented to their *unaffected* right visual space. Similarly, in recovery of motor stroke in upper limb hemiplegia, improved motor functions in the stroke-affected hand occurred following training of the unaffected hand (Manca et al., 2021).

Interestingly, our participants with mild to no leftward inattention showed increased tracking in comparison to the control condition both to the left and right of fixation. The right-sided increase was not initially hypothesized, as no such change was previously shown in neurotypical individuals (Edwards et al., 2021). In neurotypical individuals, a gain in attention was only found in the visual space opposite to where the participant attended for a prolonged duration. After isolation of attention to one side of visual space during a central fixation, we hypothesized that the ignored visual space benefited from rebalancing visual field wide attention via homeostatic gain control (Muret & Makin, 2021; Turrigiano, 2017) or post-inhibitory rebound spiking (Pugh & Raman, 2006).

However, in individuals with RCVA, we find an increase in multiple object tracking in comparison to control both in the right visual space where attention was isolated during the manipulation, and the leftward space. Similar to previous studies with chronic motor stroke (Sun et al., 2018), the individuals in our study could have experienced training in both the less-affected and more-affected visual space. However, multiple sessions of MOT are usually necessary to induce successful training in the neurotypical population (Strong & Alvarez, 2017). In patients with RCVA, sustained attention tasks also induce training over the course of days rather than minutes (Robertson et al., 1995; Van Vleet & DeGutis, 2013). Thus, while we cannot rule out a training transfer across hemifields as the cause of the bilateral attention increase, we find it unlikely with such a short (30 min) intervention.

Patients with RCVA have demonstrated nonlateralized sustained and temporal attention deficits (Posner & Peterson, 1990; Battelli et al., 2001, 2003; Husain and Rorden 2003). Control of bilateral sustained attention and temporal attention is considered right-hemisphere dominant. Training over six days using nonlateralized sustained attention tasks such as tone counting and centrally presented vertical letter cancellation has been shown to improve leftward attention (Robertson et al., 1995). Our finding may be a demonstration of how leveraging stronger rightward attentional capabilities can improve sustained attention bilaterally following isolation of attention to the right visual field. As the right visual space is less affected in patients with RCVA, there is a potential benefit of interventions performed in right visual space only. Comparably in the motor domain, less severe unilateral stroke patients with preserved corpus callosum recover motor functions better than more impaired patients. The unaffected hemisphere plays a crucial role in recovery by interhemispheric cross-facilitation of functions as evidenced by behavioral tasks (Carson 2020; Brancaccio et al., 2022). Similarly, our novel experimental protocol to promote recovery by isolating attention in the ipsilesional field might have boosted bilateral improvement in the less severe cases of leftward inattention, thanks in part to preserved structures in the left hemisphere. Interestingly, although there is previous evidence of the benefits of multiple session bilateral sustained attention training to leftward attention (Robertson et al., 1995; Van Vleet & DeGutis, 2012), we found no such benefit following our one-session bilateral control task. It is plausible that sustained attention to the unaffected visual space expedited the sustained attention improvement following just one session, rather than multiple sessions.

When correlating the impact of attention isolation on tracking with the attention characterization scores, we found a strong positive correlation between left-sided VRLAT and differences between the intervention and control tasks at the post-intervention time point: individuals who performed better in identifying objects to the left of the path in the VRLAT also had a bigger difference in tracking abilities in the intervention compared with the control. The VRLAT has previously been shown to be highly sensitive in identifying patients with leftward inattention, including those with milder symptoms which may remain undetected with paper-and-pencil tests (Dawson et al., 2008; Buxbaum et al., 2012; Corti et al., 2021). This correlation suggests that the VRLAT may be a useful tool for determining the impact of an intervention.

Although this study outlines the potential for a novel intervention, there are follow-up questions to address. First, how long does the intervention impact last, and can the length of the intervention impact be increased by multiple sessions? Although the goal of many treatments for cognitive change is to provide the longest impact of intervention over the shortest number of training sessions (Edwards et al., 2020; Contò et al., 2021), effective interventions often require multiple training days. Here, we only recorded the impact of the intervention over a ten-minute period directly after intervention. Monitoring the impact of invention over time, and then introducing multiple sessions to boost the longevity of the intervention could increase the efficacy of the treatment, potentially lasting well beyond the intervention period (Herpich et al., 2019). Second, does the intervention generalize to leftward and visual field-wide attention tasks other than the multiple object tracking task performed in this study? A critical efficacy measure of any intervention is the impact of that intervention on other tasks involving the cognitive function which has been targeted by the intervention. The generalization should be tested with other controlled attentional tasks, and in real-world tasks (Pierce & Buxbaum, 2002; Katz et al., 2005). Finally, could this intervention be useful in participants demonstrating attention deficits following traumatic brain injury (TBI)? Here we have focused the impact of our intervention on individuals with RCVA, however, there is a large body of evidence demonstrating individuals with TBI may also exhibit leftward attention related deficits (Ponsford & Kinsella, 1992; Ghajar & Ivery, 2008; Kim et al., 2008; Bonnì et al., 2012). The impact of TBI on sustained attention (Richard et al., 2018) and the effectiveness of sustained attention training in individuals with TBI (Wilson & Manly, 2003) suggests our intervention may also be successful with such individuals.

One limitation of the current study is the number of participants (n=11) included in the study (Azouvi et al., 2017). Although the study provides a within-subjects control for each participant within the study, we acknowledge that the findings are preliminary.

## Conclusion

In participants with mild to no leftward inattention, we found greater bilateral sustained attention following isolation of attention to the right of fixation when compared with a control condition. This intervention only requires 30 minutes of sustained attention in the unaffected visual space of the participant. However, we did not find a specific pre-to post-intervention impact in the participants with mild to no leftward inattention. Our attention increase is specific to sustained attention, with attention greater following the sustained attention intervention in comparison to a control sustained attention task.

## Data & Analysis Scripts

Data and analysis scripts are available to reviewers and will be available to all on OSF when published.

## Acknowledgements

Data collection funded by MossRehab Research Resource Utilization. L. Battelli is supported by the NIH (R01 AG060981-01), the Alzheimer’s Drug and Discovery Foundation (ADDF) and the Blavatnik Family Foundation. G. Chen was supported by the Intramural Research Program of the NIMH (ZICMH002888).

## Contributions

G. Edwards, L. Buxbaum, D. Edwards, & L. Battelli designed the experiment. G. Edwards performed the experiments. G. Edwards & G. Chen analyzed the data. G. Edwards, L. Buxbaum, D. Edwards, & L. Battelli wrote the manuscript.

## References

Agosta, S., Herpich, F., Miceli, G., Ferraro, F., & Battelli, L. (2014). Contralesional rTMS relieves visual extinction in chronic stroke. Neuropsychologia, 62, 269–276.

Appelros, P., Nydevik, I., Karlsson, G. M., Thorwalls, A., & Seiger, Å. (2004). Recovery from unilateral neglect after right-hemisphere stroke. Disability and Rehabilitation, 26(8), 471–477.

Azouvi, P., Jacquin-Courtois, S., & Luauté, J. (2017). Rehabilitation of unilateral neglect: Evidence-based medicine. Annals of Physical and Rehabilitation Medicine, 60(3), 191–197. https://doi.org/10.1016/j.rehab.2016.10.006

Battelli, L., Cavanagh, P., Intriligator, J., Tramo, M. J., Hénaff, M.-A., Michèl, F., & Barton, J. J. (2001). Unilateral right parietal damage leads to bilateral deficit for high-level motion. Neuron, 32(6), 985–995.

Battelli, L., Cavanagh, P., & Thornton, I. M. (2003). Perception of biological motion in parietal patients. Neuropsychologia, 41(13), 1808–1816.

Barrett, A. M., Buxbaum, L. J., Coslett, H. B., Edwards, E., Heilman, K. M., Hillis, A. E., Milberg, W. P., & Robertson, I. H. (2006). Cognitive rehabilitation interventions for neglect and related disorders: Moving from bench to bedside in stroke patients. Journal of Cognitive Neuroscience, 18(7), 1223–1236.

Bartolomeo, P., & Malkinson, T. S. (2019). Hemispheric lateralization of attention processes in the human brain. Current Opinion in Psychology, 29, 90–96.

Bates, D., Mächler, M., Bolker, B., & Walker, S. (2014). Fitting linear mixed-effects models using lme4. ArXiv Preprint ArXiv:1406.5823.

Becker, E., & Karnath, H.-O. (2007). Incidence of visual extinction after left versus right hemisphere stroke. Stroke, 38(12), 3172–3174.

Benwell, C. S. Y., Thut, G., Learmonth, G., & Harvey, M. (2013). Spatial attention: Differential shifts in pseudoneglect direction with time-on-task and initial bias support the idea of observer subtypes. Neuropsychologia, 51(13), 2747–2756. https://doi.org/10.1016/j.neuropsychologia.2013.09.030

Bisiach, E., Bulgarelli, C., Sterzi, R., & Vallar, G. (1983). Line bisection and cognitive plasticity of unilateral neglect of space. Brain and Cognition, 2(1), 32–38. https://doi.org/10.1016/0278-2626(83)90027-1

Brancaccio, A., Tabarelli, D., & Belardinelli, P. (2022). A New Framework to Interpret Individual Inter-Hemispheric Compensatory Communication after Stroke. Journal of Personalized Medicine, 12(1), 59.

Brighina, F., Bisiach, E., Oliveri, M., Piazza, A., La Bua, V., Daniele, O., & Fierro, B. (2003). 1 Hz repetitive transcranial magnetic stimulation of the unaffected hemisphere ameliorates contralesional visuospatial neglect in humans. Neuroscience Letters, 336(2), 131–133.

Bonnì, S., Mastropasqua, C., Bozzali, M., Caltagirone, C., & Koch, G. (2013). Theta burst stimulation improves visuo-spatial attention in a patient with traumatic brain injury. Neurological Sciences, 34(11), 2053–2056.

Buxbaum LJ, Dawson AM, Linsley D. Reliability and validity of the Virtual Reality Lateralized Attention Test in assessing hemispatial neglect in right-hemisphere stroke. Neuropsychology. 2012 Jul;26(4):430–41.

Carson, R. G. (2020). Inter-hemispheric inhibition sculpts the output of neural circuits by co-opting the two cerebral hemispheres. The Journal of Physiology, 598(21), 4781–4802.

Cavézian, C., Danckert, J., Lerond, J., Daléry, J., d’Amato, T., & Saoud, M. (2007). Visual-perceptual abilities in healthy controls, depressed patients, and schizophrenia patients. Brain and Cognition, 64(3), 257–264. https://doi.org/10.1016/j.bandc.2007.03.008

Contò, F., Edwards, G., Tyler, S., Parrott, D., Grossman, E., & Battelli, L. (2021). Attention network modulation via tRNS correlates with attention gain. Elife, 10, e63782.

Corbetta, M., Kincade, M. J., Lewis, C., Snyder, A. Z., & Sapir, A. (2005). Neural basis and recovery of spatial attention deficits in spatial neglect. Nature Neuroscience, 8(11), 1603–1610.

Corbetta, M., & Shulman, G. L. (2002). Control of goal-directed and stimulus-driven attention in the brain. Nature Reviews Neuroscience, 3(3), 201–215.

Corti, C., Oprandi, M. C., Chevignard, M., Jansari, A., Oldrati, V., Ferrari, E., Martignoni, M., Romaniello, R., Strazzer, S., & Bardoni, A. (2021). Virtual-Reality Performance-Based Assessment of Cognitive Functions in Adult Patients With Acquired Brain Injury: A Scoping Review. Neuropsychology Review. https://doi.org/10.1007/s11065-021-09498-0

Culham, J. C., Brandt, S. A., Cavanagh, P., Kanwisher, N. G., Dale, A. M., & Tootell, R. B. (1998). Cortical fMRI activation produced by attentive tracking of moving targets. Journal of Neurophysiology.

Dawson, A. M., Buxbaum, L. J., & Rizzo, A. A. (2008). The virtual reality lateralized attention test: Sensitivity and validity of a new clinical tool for assessing hemispatial neglect. 2008 Virtual Rehabilitation, 77–82.

Edwards, G., Berestova, A., & Battelli, L. (2021). Behavioral gain following isolation of attention. Scientific Reports, 11(1), 1–10.

Frassinetti, F., Angeli, V., Meneghello, F., Avanzi, S., & Làdavas, E. (2002). Long-lasting amelioration of visuospatial neglect by prism adaptation. Brain, 125(3), 608–623.

Gauthier, L., Dehaut, F., & Joanette, Y. (1989). The bells test: A quantitative and qualitative test for visual neglect. International Journal of Clinical Neuropsychology, 11(2), 49–54.

Ghajar, J., & Ivry, R. B. (2008). The predictive brain state: Timing deficiency in traumatic brain injury? Neurorehabilitation and Neural Repair, 22(3), 217–227.

Gillen, R., Tennen, H., & McKee, T. (2005). Unilateral spatial neglect: Relation to rehabilitation outcomes in patients with right hemisphere stroke. Archives of Physical Medicine and Rehabilitation, 86(4), 763–767.

Herpich, F., Melnick, M. D., Agosta, S., Huxlin, K. R., Tadin, D., & Battelli, L. (2019). Boosting learning efficacy with noninvasive brain stimulation in intact and brain-damaged humans. Journal of Neuroscience, 39(28), 5551–5561.

Husain, M., & Rorden, C. (2003). Non-spatially lateralized mechanisms in hemispatial neglect. Nature Reviews Neuroscience, 4(1), 26–36.

Katz, N., Ring, H., Naveh, Y., Kizony, R., Feintuch, U., & Weiss, P. L. (2005). Interactive virtual environment training for safe street crossing of right hemisphere stroke patients with Unilateral Spatial Neglect. Disability and Rehabilitation, 27(20), 1235–1244. https://doi.org/10.1080/09638280500076079

Kim, Y.-H., Yoo, W.-K., Ko, M.-H., Park, C., Kim, S. T., & Na, D. L. (2009). Plasticity of the Attentional Network After Brain Injury and Cognitive Rehabilitation. Neurorehabilitation and Neural Repair, 23(5), 468–477. https://doi.org/10.1177/1545968308328728

Kinsbourne, M. (1987). Mechanisms of unilateral neglect. In Advances in psychology (Vol. 45, pp. 69–86). Elsevier.

Kinsbourne, M. (1993). Orientational bias model of unilateral neglect: Evidence from attentional gradients within hemispace. Í Robertson IH, Marshall J ritstj. Unilateral neglect: Clinical and Experimental studies. Unilateral Neglect: Clinical and Experimental Studies. Hove, UK: Lawrence Erlbaum;, 63–86.

Kinsbourne, M. (1994). Mechanisms of neglect: Implications for rehabilitation. Neuropsychological Rehabilitation, 4(2), 151–153.

Làdavas, E., Menghini, G., & Umiltà, C. (1994). A rehabilitation study of hemispatial neglect. Cognitive Neuropsychology, 11(1), 75–95. https://doi.org/10.1080/02643299408251967

Learmonth, G., Märker, G., McBride, N., Pellinen, P., & Harvey, M. (2018). Right-lateralised lane keeping in young and older British drivers. PLOS ONE, 13(9), e0203549. https://doi.org/10.1371/journal.pone.0203549

Learmonth, G., & Papadatou-Pastou, M. (2021). A Meta-Analysis of Line Bisection and Landmark Task Performance in Older Adults. Neuropsychology Review. https://doi.org/10.1007/s11065-021-09505-4

Lenth, R., Singmann, H., Love, J., Buerkner, P., & Herve, M. (2018). Emmeans: Estimated marginal means, aka least-squares means. R Package Version, 1(1), 3.

Lindsay, M. P., Norrving, B., Sacco, R. L., Brainin, M., Hacke, W., Martins, S., Pandian, J., & Feigin, V. (2019). World Stroke Organization (WSO): Global stroke fact sheet 2019. SAGE Publications Sage UK: London, England.

Manca, A., Hortobágyi, T., Carroll, T. J., Enoka, R. M., Farthing, J. P., Gandevia, S. C., Kidgell, D. J., Taylor, J. L., & Deriu, F. (2021). Contralateral effects of unilateral strength and skill training: Modified delphi consensus to establish key aspects of cross-education. Sports Medicine, 51(1), 11–20.

Martinez, V., & Sarter, M. (2004). Lateralized attentional functions of cortical cholinergic inputs. Behavioral Neuroscience, 118(5), 984.

Muret, D., & Makin, T. R. (2021). The homeostatic homunculus: Rethinking deprivation-triggered reorganisation. Current Opinion in Neurobiology, 67, 115–122.

Ogden, J. A. (1985). Anterior-posterior interhemispheric differences in the loci of lesions producing visual hemineglect. Brain and Cognition, 4(1), 59–75.

Parton, A., Malhotra, P., & Husain, M. (2004). Hemispatial neglect. Journal of Neurology, Neurosurgery & Psychiatry, 75(1), 13–21.

Pierce, S. R., & Buxbaum, L. J. (2002). Treatments of unilateral neglect: A review. Archives of Physical Medicine and Rehabilitation, 83(2), 256–268. https://doi.org/10.1053/apmr.2002.27333

Ponsford, J., & Kinsella, G. (1992). Attentional deficits following closed-head injury. Journal of Clinical and Experimental Neuropsychology, 14(5), 822–838. https://doi.org/10.1080/01688639208402865

Posner, M. I., & Petersen, S. E. (1990). The attention system of the human brain. Annual Review of Neuroscience, 13(1), 25–42.

Pugh, J. R., & Raman, I. M. (2006). Potentiation of mossy fiber EPSCs in the cerebellar nuclei by NMDA receptor activation followed by postinhibitory rebound current. Neuron, 51(1), 113–123.

Pylyshyn, Z. W., & Storm, R. W. (1988). Tracking multiple independent targets: Evidence for a parallel tracking mechanism. Spatial Vision, 3(3), 179–197.

R Core Team (2020). —European Environment Agency. (n.d.). [Methodology Reference]. Retrieved January 5, 2022, from https://www.eea.europa.eu/data-and-maps/indicators/oxygen-consuming-substances-in-rivers/r-development-core-team-2006

Richard, N. M., O’Connor, C., Dey, A., Robertson, I. H., & Levine, B. (2018). External modulation of the sustained attention network in traumatic brain injury. Neuropsychology, 32(5), 541.

Robertson, I. H., Tegnér, R., Tham, K., Lo, A., & Nimmo-smith, I. (1995). Sustained attention training for unilateral neglect: Theoretical and rehabilitation implications. Journal of Clinical and Experimental Neuropsychology, 17(3), 416–430. https://doi.org/10.1080/01688639508405133

Rossetti, Y., Rode, G., Pisella, L., Farné, A., Li, L., Boisson, D., & Perenin, M.-T. (1998). Prism adaptation to a rightward optical deviation rehabilitates left hemispatial neglect. Nature, 395(6698), 166–169.

Schenkenberg, T., Bradford, D. C., & Ajax, E. T. (1980). Line bisection and unilateral visual neglect in patients with neurologic impairment. Neurology, 30(5), 509–509.

Shipp, S. (2004). The brain circuitry of attention. Trends in Cognitive Sciences, 8(5), 223–230.

Silasi, G., & Murphy, T. H. (2014). Stroke and the connectome: How connectivity guides therapeutic intervention. Neuron, 83(6), 1354–1368.

Strong, R. W., & Alvarez, G. A. (2017). Training enhances attentional expertise, but not attentional capacity: Evidence from content-specific training benefits. Journal of Vision, 17(4), 4–4.

Sun, Y., Ledwell, N. M., Boyd, L. A., & Zehr, E. P. (2018). Unilateral wrist extension training after stroke improves strength and neural plasticity in both arms. Experimental Brain Research, 236(7), 2009–2021.

Turrigiano, G. G. (2017). The dialectic of Hebb and homeostasis. Philosophical Transactions of the Royal Society B: Biological Sciences, 372(1715), 20160258.

Van Vleet, T. M., & DeGutis, J. M. (2013). Cross-training in hemispatial neglect: Auditory sustained attention training ameliorates visual attention deficits. Cortex, 49(3), 679–690. https://doi.org/10.1016/j.cortex.2012.03.020

Wilson, B., Cockburn, J., & Halligan, P. (1987). Development of a behavioral test of visuospatial neglect. Archives of Physical Medicine and Rehabilitation, 68(2), 98– 102.

Winstein, C. J., Stein, J., Arena, R., Bates, B., Cherney, L. R., Cramer, S. C., Deruyter, F., Eng, J. J., Fisher, B., & Harvey, R. L. (2016). Guidelines for adult stroke rehabilitation and recovery: A guideline for healthcare professionals from the American Heart Association/American Stroke Association. Stroke, 47(6), e98– e169.

Yantis, S. (1992). Multielement visual tracking: Attention and perceptual organization. Cognitive Psychology, 24(3), 295–340.

Zeltzer, L., Menon, A., Korner-Bitensky, N., & Sitcoff, E. (2008). Line bisection test. Stroke Engine. Retrieved July, 18, 2018.

