## Supplemental Information for "Improving visual attention following right hemisphere stroke: A preliminary study"

Appendix 1 - Attention Characterization Tasks:

#### Line Bisection Task

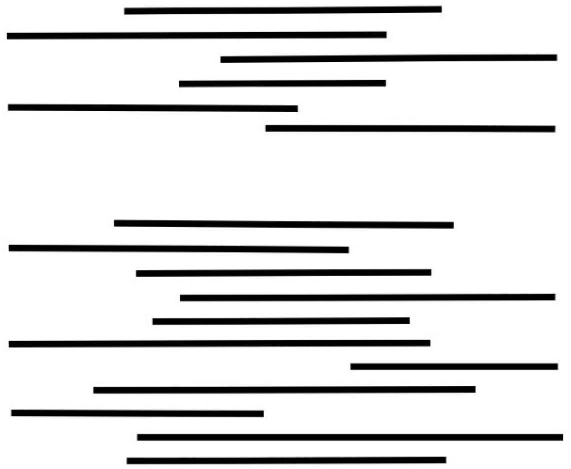

#### Bells Test

Task:

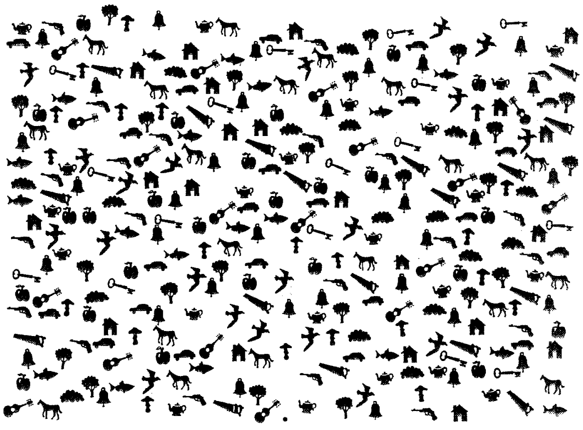

Score Sheet:

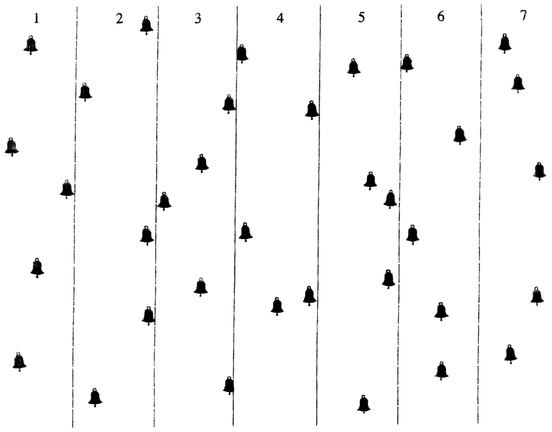

### Virtual Reality Lateralized Attention Test (VRLAT) Score Sheet

**SCORESHEET**

**VIRTUAL REALITY LATERALIZED ATTENTION TEST (VRLAT)**

**Laurel J. Buxbaum, Dean Klimchuk, & Roman Mitura**

Moss Rehabilitation Research Institute, Elkins Park, PA, and Digital Mediaworks, Ontario, CA

**PARTICIPANT**________________________________ **DATE**________________

**ARRAY TYPE(circle):** **Simple** **Complex** **Enhanced**

**DRIVER (circle):** **Participant** **Examiner**

**(circle detected targets below)**

| **FORWARD** | | | **REVERSED** | | |
|---|---|---|---|---|---|
| TARGET | LOCATION | NOTES | TARGET | LOCATION | NOTES |
| orange tree | L | | red tree | R | |
| horse | R | | lion | L | |
| pig | L | | purple tree | L | |
| white tree | R | | elephant | L | |
| turtle | R | | blue tree | R | |
| camel | L | | dolphin | R | |
| green tree | R | | yellow tree | R | |
| cat/lion | L | | pink tree | L | |
| black tree | R | | dog | R | |
| cow | R | | brown tree | R | |
| brown tree | L | | cow | L | |
| dog | L | | black tree | L | |
| pink tree | R | | cat/lion | R | |
| yellow tree | L | | green tree | L | |
| dolphin | L | | camel | R | |
| blue tree | L | | turtle | L | |
| elephant | R | | white tree | L | |
| purple tree | R | | pig | R | |
| lion | R | | horse | L | |
| red tree | L | | orange tree | R | |

TOTAL FORWARD________________ TOTAL REVERSED ________________

GRAND TOTAL______________

Full Scoring:
0 = nothing
1 = 'something'
2 = category or color error
3 = complete identification

Simple Scoring: Award 1 point for each detection

©Copyright Einstein Healthcare Network 2013

### Supplementary Information

#### Eyetracking Data

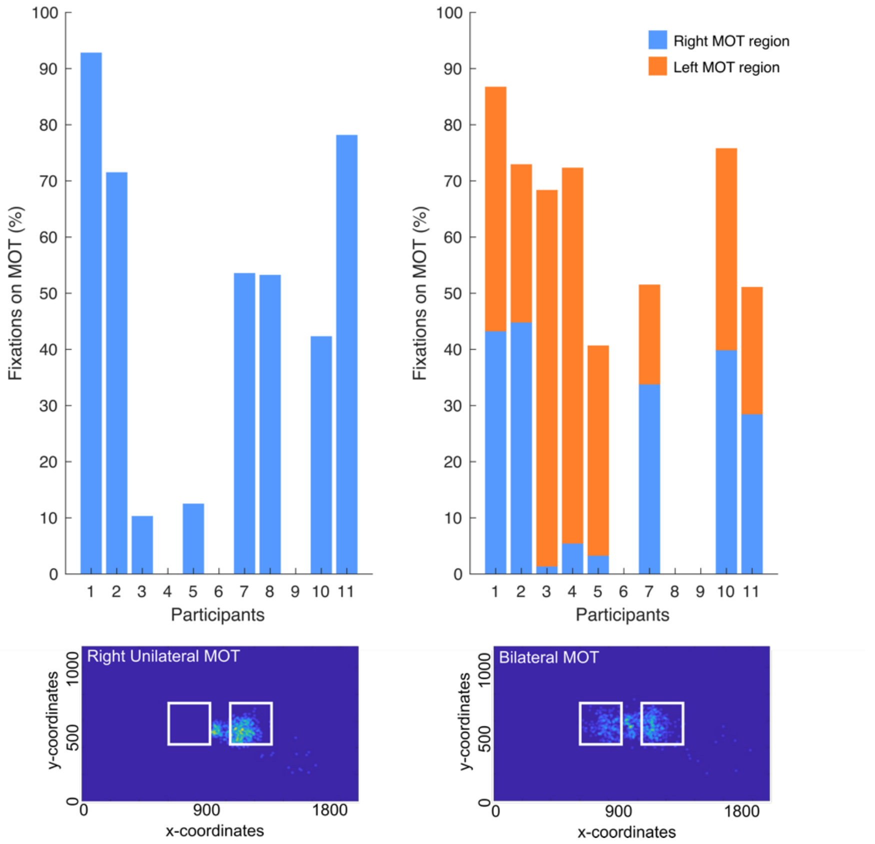


**Supplemental Figure 1: Percentage of fixations on unilateral and bilateral multiple object tracking stimuli.** Top right panel denotes number of fixations on right unilateral MOT during the intervention session. Bottom right panel: example illustration of fixations from one participant during right unilateral MOT. Top left panel illustrates the number of fixations on right and left MOT during bilateral MOT control. Bottom left panel: example illustration of fixations from one participant during bilateral MOT.

#### Model Comparisons

Model comparisons were conducted on various *glmer* models to determine the best model fit for the data collected.

**Key:**
**Accuracy = Acc; Session = Sess; Manip = manipulation VF = Visual field/ space; Inatt = inattention score; ET = eye-tracking – the percentage fixations in right visual space; Subs = participants**

**Syntax:**

Mod1 <- glmer (Acc ~ Sess + VF + Manip + Inatt + ET + *(1+Sess*|*Subs)* + *(1+VF*|*Subs)* + *(1*+ Manip|*Subs),* data = AttIso, family = binomial)

Mod2 <- glmer (Acc ~ Sess * VF + Manip + Inatt + ET + *(1+ Sess |Subs)* + *(1+VF|Subs)* + *(1+* Manip|*Subs),* data = AttIso, family = binomial)
Mod3 <- glmer (Acc ~ Sess + VF * Manip + Inatt + ET + *(1+ Sess |Subs)* + *(1+VF|Subs)* + *(1+* Manip|*Subs),* data = AttIso, family = binomial)
Mod4 <- glmer (Acc ~ Sess + VF + Manip * Inatt + ET *(1+ Sess |Subs)* + *(1+VF|Subs)* + *(1+* Manip|*Subs),* data = AttIso, family = binomial)
Mod5 <- glmer (Acc ~ Sess + VF * Inatt + Manip + ET + *(1+ Sess |Subs)* + *(1+VF|Subs)* + *(1+* Manip|*Subs),* data = AttIso, family = binomial)
Mod6 <- glmer (Acc ~ Sess * Inatt + Manip + VF + ET + *(1+ Sess |Subs)* + *(1+VF|Subs)* + *(1+* Manip|*Subs),* data = AttIso, family = binomial)
Mod7 <- glmer (Acc ~ Sess * Manip + Inatt + VF + ET + *(1+ Sess |Subs)* + *(1+VF|Subs)* + *(1+* Manip|*Subs),* data = AttIso, family = binomial)
Mod8 <- glmer (Acc ~ Sess * Manip * Inatt + VF + ET + *(1+ Sess |Subs)* + *(1+VF|Subs)* + *(1+* Manip|*Subs),* data = AttIso, family = binomial)
Mod9 <- glmer (Acc ~ Sess + Manip * Inatt * VF + ET + *(1+ Sess |Subs)* + *(1+VF|Subs)* + *(1+* Manip|*Subs),* data = AttIso, family = binomial)
Mod10 <- glmer (Acc ~ Sess * Manip + Inatt * VF + ET + *(1+ Sess |Subs)* + *(1+VF|Subs)* + *(1+* Manip|*Subs),* data = AttIso, family = binomial)
Mod11 <- glmer (Acc ~ Sess * VF + Manip * Inatt + ET + *(1+ Sess |Subs)* + *(1+VF|Subs)* + *(1+* Manip|*Subs),* data = AttIso, family = binomial)
Mod12 <- glmer (Acc ~ Sess * VF * Manip * Inatt + ET + *(1+ Sess |Subs)* + *(1+VF|Subs)* + *(1+* Manip|*Subs),* data = AttIso, family = binomial)

anova(Mod1, Mod2, Mod3, Mod4, Mod5, Mod6, Mod7, Mod8, Mod9, Mod10, Mod11, Mod12)

| Mod | npar | AIC | BIC | Loglik | deviance | Chisq | Chi Df | Pr(>Chisq) |
|---|---|---|---|---|---|---|---|---|
| Mod1 | 15 | 2567.0 | 2650.8 | -1268.5 | 2537.0 | | | |
| Mod3 | 16 | 2569.0 | 2658.4 | -1268.5 | 2537.0 | 0.0073 | 1 | 0.93203 |
| Mod2 | 16 | 2567.1 | 2656.4 | -1267.5 | 2535.1 | 1.9579 | 0 | |
| Mod4 | 16 | 2565.2 | 2654.5 | -1266.6 | 2533.2 | 1.9143 | 0 | |
| Mod5 | 16 | 2568.9 | 2658.3 | -1268.5 | 2536.9 | 0.000 | 0 | |
| Mod6 | 16 | 2568.8 | 2658.2 | -1268.4 | 2536.8 | 0.1171 | 0 | |
| Mod7 | 16 | 2568.9 | 2658.2 | -1268.4 | 2536.9 | 0.000 | 0 | |
| Mod10 | 17 | 2570.8 | 2665.7 | -1268.4 | 2536.8 | 0.0994 | 1 | 0.75256 |
| Mod11 | 17 | 2567.2 | 2662.1 | -1266.6 | 2533.2 | 3.6261 | 0 | |
| Mod8 | 19 | 2563.3 | 2669.4 | -1262.7 | 2525.3 | 7.8318 | 2 | 0.01992 |
| Mod9 | 19 | 2568.2 | 2674.3 | -1265.1 | 2530.2 | 0.000 | 0 | |
| Mod12 | 26 | 2574.0 | 2719.2 | -1261.0 | 2522.0 | 8.2168 | 7 | 0.31387 |

Model comparisons demonstrated Mod8 is the least complex and best fit for our data.
